# A bioreactor for controlled electromechanical stimulation of developing scaffold-free constructs

**DOI:** 10.1101/2021.01.10.426136

**Authors:** Sarah K. Van Houten, Michael T. K. Bramson, David T. Corr

## Abstract

Bioreactors are commonly used to apply biophysically-relevant stimulations to tissue-engineered constructs in order to explore how these stimuli influence tissue development, healing, and homeostasis. These bioreactors offer great flexibility as key features of the stimuli (*e*.*g*., duty cycle, frequency, amplitude, duration) can be controlled to elicit a desired cellular response. Controlled delivery of mechanical and/or electrical stimulation has been shown to improve the structure and function of engineered tissue constructs, compared to unstimulated controls, for applications in various musculoskeletal soft tissues. However, most bioreactors that apply mechanical and electrical stimulations, do so to a scaffold after the construct has developed, preventing study of the influence these stimuli have on early construct development. Thus, there is a need for a bioreactor that allows the direct application of mechanical and electrical stimulation to constructs **as they develop**, to enable such exploration and to better mimic key aspects of tissue development. Hence, the objective of this study was to develop and calibrate a bioreactor to deliver precise mechanical and electrical stimulation, either independently or in combination, to developing scaffold-free tissue constructs. Standard Flexcell Tissue Train plates were modified with stainless steel loading pins and stimulating electrodes to integrate direct mechanical and electrical stimulation, respectively, into our established scaffold-free, single-fiber engineering platform. Linear calibration curves were established, then used to apply precise dynamic mechanical and electrical stimulations, over a range of physiologically relevant strains (0.50, 0.70, 0.75, 1.0, 1.5%) and voltages (1.5, 3.5 V), respectively. Once calibrated, applied mechanical and electrical stimulations were not statistically different from their desired target values, and were consistent whether delivered independently or in combination. Indeed, concurrent delivery of mechanical and electrical stimulation resulted in a negligible change in mechanical (< 2%) and electrical (<1%) values, from their independently-delivered values. With this calibrated bioreactor, we can apply precise, controlled, reproducible mechanical and electrical stimulations, alone or in combination, to scaffold-free, tissue engineered constructs as they develop, to explore how these stimuli can be leveraged to advance and accelerate functional tissue engineering in a variety of musculoskeletal soft tissues.

**IMPACT STATEMENT:** As the importance of biophysical stimulation in tissue engineering continues to be recognized and incorporated, this bioreactor provides a platform to further our understanding of the roles independent or combined mechanical and electrical stimulation have in tissue development and functional maturation, and may inform future tissue engineering approaches for clinical applications.

## 1. INTRODUCTION

In recent years, tissue engineers have sought to assess the impacts of microenvironmental stimuli on tissue development and function, aiming to overcome financial and functional shortcomings of conventional treatments (*i*.*e*., surgical replacement grafts).^1,2^ Despite many notable advances toward engineering clinically implantable tissue constructs,^1–3^ creating scalable constructs that replicate the structure and function of native tissue remains a challenge. One approach tissue engineers use to address this challenge is to deliver physiologic-like stimulation (*e*.*g*., mechanical, electrical) to engineered constructs using custom-built bioreactors.^4–6^ Bioreactors offer great flexibility because the stimuli’s key features (*e*.*g*., duty cycle, frequency, amplitude, duration, waveform) can be controlled to elicit a desired response. By mirroring complex stimulation patterns seen *in vivo* during development and healing, tissue engineers can leverage corresponding biophysical responses to accelerate construct maturation and tune the structure and biomechanical properties of engineered tissue constructs.

Tissue engineers can apply mechanical stimulation using various methods including spinner flasks (shear forces),^7^ compressive pistons,^7^ or step-motors which allow for uniaxial strain loading and are used in many different tissue engineering platforms (*e*.*g*., tendon, ligament, skeletal muscle).^3,8^ Additionally, cell culture systems, such as the Flexcell system, have been widely used to deliver dynamic mechanical stimulation patterns to deformable substrates through the application of computer-controlled vacuum pressure.^9^ Mechanical loading is traditionally applied to a scaffold-based structure in which cells are seeded, or to engineered constructs after their cells have laid down sufficient extracellular matrix to produce tissue structure. Incorporation of mechanical stimulation in tissue engineering approaches has been shown to improve construct alignment, increase construct thickness and cell matrix production, and promote cell elongation and fusion.^10–12^

Electrical stimulation has also been explored for engineering various musculoskeletal tissues (*e*.*g*., skeletal muscle, cardiac, bone).^13–15^ Stimuli are applied via electric field or direct stimulation, and while the method of stimulation may vary, the primary goal is typically to mimic the neural stimulation cells experience *in vivo*. Electrical stimulation has been shown to increase construct alignment and cell proliferation, and promote differentiation and growth factor production.^16,17^ In addition to these structural and compositional benefits, electrical stimulation has been shown to activate different cell types in bone, and generate active force production in cardiac and skeletal muscle tissues.^13–16^

Due to the demonstrated individual benefits of electrical stimulation and mechanical strain, and considering that these stimuli often occur simultaneously within the body, a number of recent studies have sought to explore potential synergistic benefits of concurrent mechanical and electrical stimulation.^17–20^ In engineered skeletal muscle, combined stimulation increased the amount of fast myosin heavy chain proteins present,^17^ and increased construct thickness and force production, when compared to unstimulated controls.^17–20^ These promising findings hint that such synergies exist, and may factor prominently in future tissue engineering strategies. Hence, there is a need for more studies that explore the possible synergistic effects of concurrent mechanical and electrical stimulation on tissue engineered constructs, as well as the effects these stimuli have on construct development and subsequent biomechanical function.^6,20,21^

The vast majority of tissue engineering approaches utilize either natural or synthetic material scaffolds as a provisional structure into which cells are seeded, and can deposit their own extracellular matrix.^22^ While scaffold-based strategies offer advantages such as design flexibility and structural support,^23^ they may limit direct stimulation of cells within the construct due to stress shielding by the scaffold, which could attenuate the cellular response.^23^ This has led some to explore scaffold-free approaches, which rely on cellular self-assembly and mechanobiological matrix production to create tissue constructs, and allow for direct stimulation of cells within them.^24,25^ However, scaffold-free approaches face technical challenges in delivering biophysical stimuli to a construct without significant initial structure. Thus, there is a need for a bioreactor capable of applying precise mechanical and electrical stimulation to scaffold-free constructs, as they develop, to enable investigation into the early effects of biophysical stimulation in tissue development and functional maturation.

Our laboratory established a scaffold-free, single-fiber tissue engineering platform, wherein cellular fibers form via directed self-assembly, with no provisional matrix present.^26,27^ We utilized this platform to engineer single tendon fibers, and applied cyclic, low-amplitude uniaxial tensile strain during fiber development to promote matrix production and functional maturation, using fiber growth channel assemblies coupled to a modified Flexcell system.^26,27^ The addition of mechanical stimulation dramatically increased fiber longevity, tensile strength, Young’s modulus, and toughness, compared to unstimulated controls.^27^ To expand this for engineering active tissue (*i*.*e*., skeletal muscle), we now seek to incorporate electrical stimulation into this platform. In this study, we integrate the ability to apply precise, custom electrical stimulation, during fiber development, into our scaffold-free single fiber engineering platform, and perform robust calibration of mechanical and electrical stimuli, delivered independently and concurrently. We then characterize the applied calibrated signals to ensure controlled, reproducible, and accurate cellular stimulation. Successful delivery and characterization of the applied stimuli to constructs as they develop will provide novel insight on the roles of mechanical and electrical stimulation in tissue development and functional maturation, and how these stimuli might be used to further tissue engineering approaches.

## 2. METHODS

### 2.1. System Overview

In this bioreactor design, we utilize our established platform for scaffold-free, single fiber engineering, wherein cells (*e*.*g*., human dermal fibroblasts, myoblasts) - densely seeded in a fibronectin-coated agarose growth channel with no provisional matrix - self-assemble to form mechanically-testable musculoskeletal fibers.^26–29^ Collagen disks at either end of the growth channel serve as anchor points for the developing fiber. As the cells align, form cell-cell junctions and self-assemble into a fiber, they develop self-generated static tension and the fiber detaches from the growth channel. The result is a single fiber spanning the entire length of the channel, suspended between the collagen disks. In our previous work, we modified Flexcell Tissue Train plate wells to include vertical stainless steel loading pins, which enabled coupling of growth channel assemblies to the Flexcell system for direct mechanical stimulation of the developing fiber constructs.^26,27^ Herein, we created Dual Stimulation plates in which the stainless steel loading pins also served as stimulating electrodes, to enable delivery of electrical stimulation to the growth channel, alone or in combination with mechanical stimulation (**Fig. 1**).

**Figure 1.**
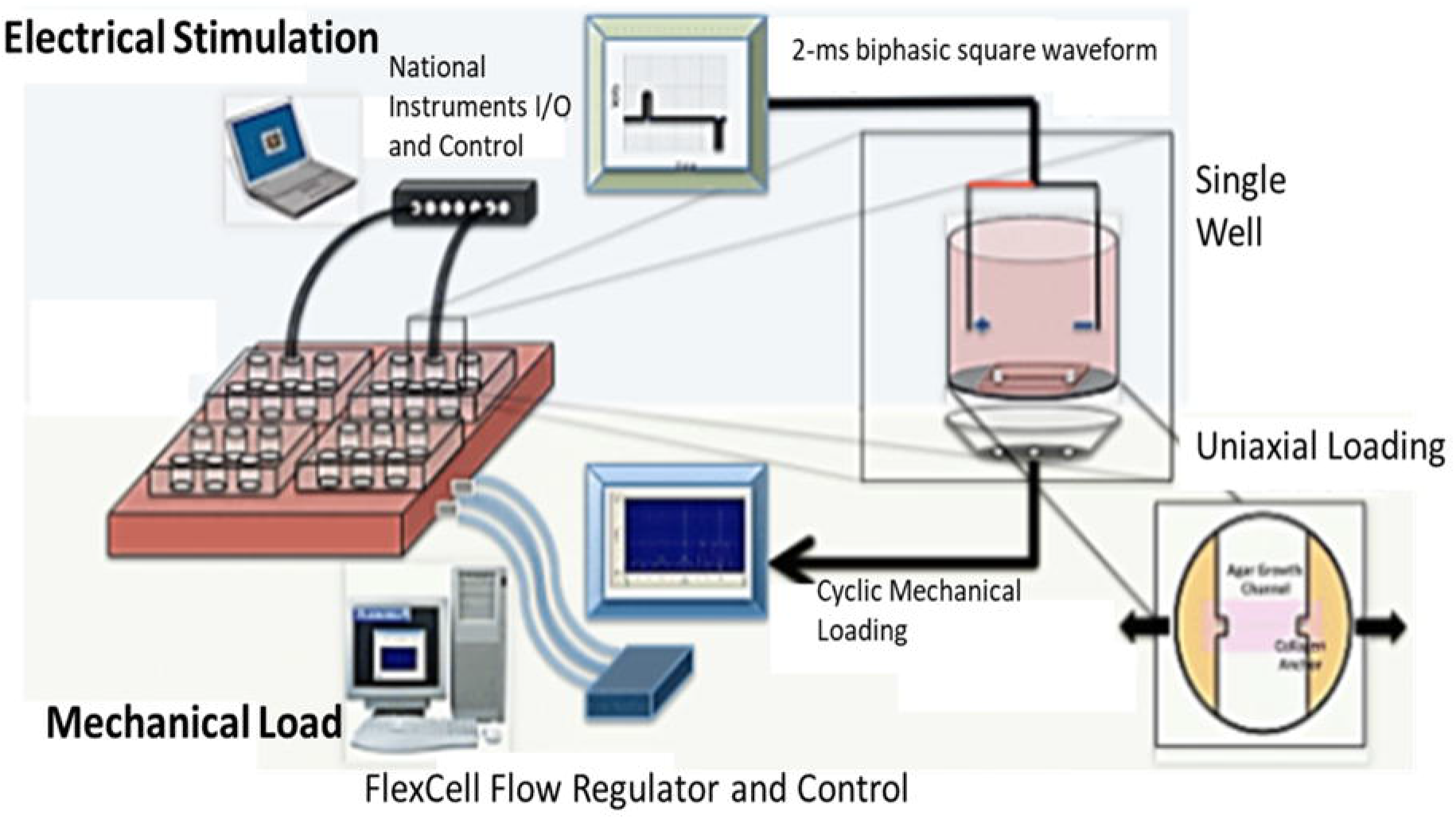
Schematic representation of a bioreactor for independent or concurrent mechanical and electrical stimulation. Electrical stimulation, programmed in LabVIEW, is applied by a National Instruments DAQ module via wires soldered to modified Flexcell plates. Mechanical loading, programmed within the Flexcell system, is applied by computer-regulated vacuum deformation of flexible multi-well plate membranes, seated in a 4-plate baseplate, allowing for simultaneous strain of up to 24 growth channel assemblies. (modified from [28]).

### 2.2. Dual Stimulation Plate Preparation

Flexcell® Tissue Train® plates (Flexcell International Corp., Burlington, NC) were modified with vertical loading pins in each of the plate’s six wells, as previously described.^27^ Briefly, 17-gauge stainless steel nails (Hillman, d = 1 mm), cut to ∼7-mm height, were attached to the nylon loading tabs of Tissue Train® plates using cyanoacrylate glue (Krazy Glue Maximum Bond Gel). To incorporate electrical stimulation, 30-gauge stainless steel wires were soldered to the loading pins, such that the pins that enable mechanical loading also function as stimulating electrodes. Soldered wires were passed out through the side of modified plates and secured in place with some slack in the wire, to provide strain relief and improve durability for repeated cyclic loading. Rectangular reinforcement tabs, fashioned out of recycled silicone Flexcell membranes and biopsy punched with 1-mm through holes, were placed over loading pins/stimulating electrodes and secured with cyanoacrylate glue to stabilize the loading pins. Once the glue had cured, the wells were rinsed with three 5-minute washes of 70% ethanol before use in experiments.

### 2.3. Growth Channel Fabrication & Use in Single-Fiber Engineering

Agarose single-fiber growth channels were fabricated using previously established methods.^27^ Briefly, a 2-wt% agarose solution was prepared by dissolving UltraPure™ Agarose (Invitrogen™, Thermo Fischer Scientific, Waltham, MA) in Dulbecco’s Modified Eagle Medium (DMEM, VWR, Bridgeport, NJ). Molten agar was then poured under a custom-fabricated aluminum 6061-T6 micro-mold (Precision MicroFab, Curtis Bay, MD) with a positive mold of the growth channel features (150-mm wide, 300-mm deep, and 17.5-mm long), inverted on 2-mm risers. After allowing the agarose to solidify for 10 minutes, the aluminum mold was lifted, leaving an exact channel with specific features in the agarose. Molded agarose growth channels were then UV-crosslinked (Spectroline, Westbury, NY) at 1 J/cm^2^ and stored at 4°C, for use within one month of fabrication.

To make growth channel assemblies for use in single-fiber engineering, sterile collagen disk anchors (d = 4 mm) punched from collagen sponge pads (Bovine Type I; Royal DSM, Exton, PA) and wet with 10 µL cell culture-grade water (Mediatech, Manassas, VA) are inserted into 4-mm holes punched at both ends of the growth channel. To couple growth channel assemblies to modified Dual Stimulation plates for stimulation, 1-mm through-holes are punched into the center of the collagen disk anchors, allowing them to be mounted onto loading pins/stimulating electrodes.^27^

Although fibers were not created as part of this study, to engineer fibers, human plasma-derived fibronectin (Corning, MA), diluted to 0.375 µg/mL, is wicked through the collagen disks into the growth channels (20 µL/channel) and allowed to dry, promoting initial cell adhesion. Cells (*e*.*g*., fibroblasts, myoblasts), cultured in standard culture conditions, are trypsinized, centrifuged, and resuspended at a high density (∼5 million cells/mL (myoblast)^30^ or 15 million cells/mL (fibroblasts)^27^) in growth medium tailored to the cell type. Cells are then seeded into the growth channel by pipetting 10 µL through each collagen disk, followed by 10 µL directly over the growth channel in each direction. Cells are allowed 10 minutes to adhere to the growth channel before immersion in growth medium and standard cell culture conditions (37°C, 5% CO_2_, and 95% relative humidity),^27^ before further stimulation is applied as early as 3 hours (myoblasts) or 18 hours (fibroblasts) post seeding.^30^

## 3. EXPERIMENT

### 3.1. EXPERIMENTAL METHODS

#### 3.1.1. Mechanical Characterization

Mechanical strain within the growth channel, indicative of the strains that cells in developing fibers would experience, was determined using Digital Image Correlation (DIC), following previously published methods.^28^ To prepare the growth channels for optical strain measurements via DIC, the top surface of each growth channel assembly was blotted dry and coated with an anisotropic, high-contrast speckle pattern of black spray paint (Rust-Oleum).^28^ 2 mL of DMEM were then added to cover the bottom of each well, mimicking typical culture conditions. Dual Stimulation plates with coupled growth channel assemblies were then mounted in a pneumatic Flexcell FX-4000 system equipped with 24-mm Arctangle loading posts (Flexcell International Corp., Burlington, NC) to enable application of controlled uniaxial strain via applied vacuum pressure (**Fig. 2**).

**Figure 2.**
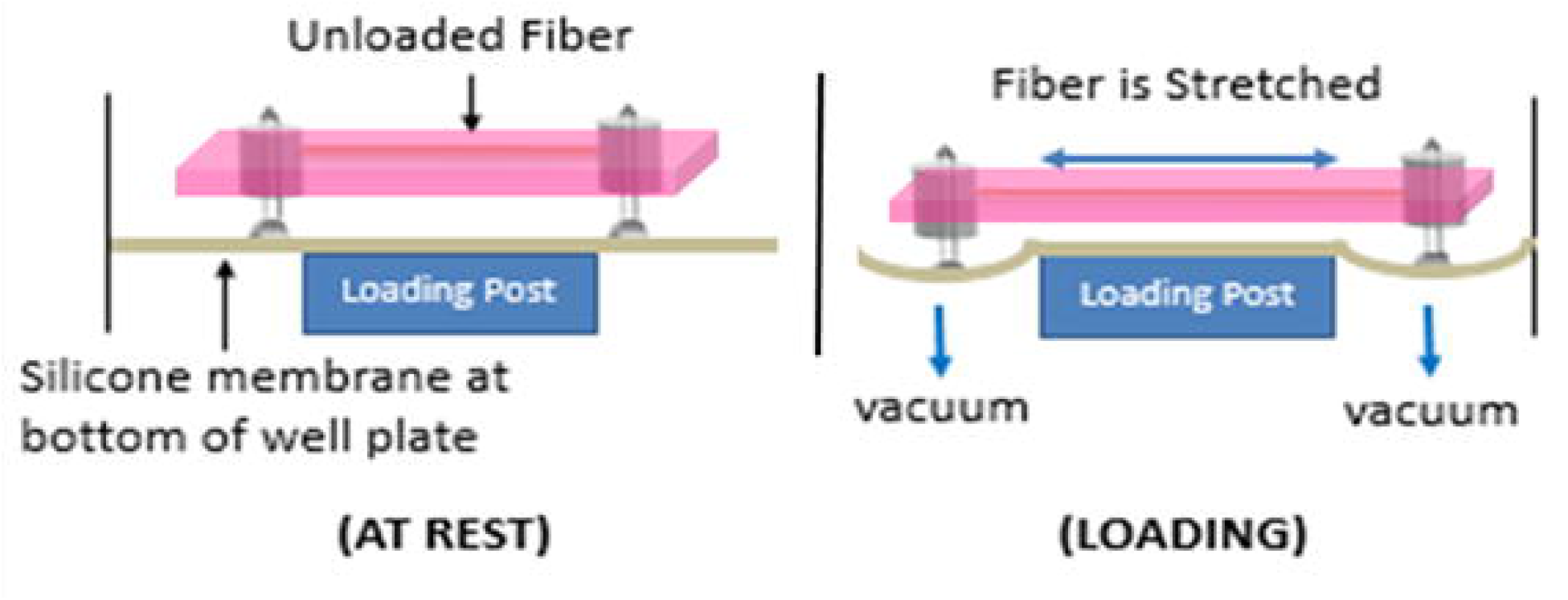
Flexcell Tissue Train plates modified with vertical loading pins allow for application of cyclic uniaxial tension to developing tissue constructs in a scaffold-free single fiber engineering approach. The Flexcell system applies user-specified strains via vacuum pressure, to uniaxially stretch the silicone membrane, driving the loading pins apart and straining the fiber growth channel (modified from [26]).

Mechanical stimulation was calibrated over a range of physiologically relevant strains, based on magnitudes and frequencies seen in musculoskeletal development.^27^ Two cycles at each of five different strain amplitudes (0.50%, 0.70%, 0.75%, 1.0%, and 1.5% strain) were applied to six independent growth channel assemblies, at three different frequencies (0.1, 0.5, and 1 Hz). A two-camera Digital Image Correlation system (Correlated Solutions, Inc., Columbia, SC) was used to dynamically measure the strain within the growth channels, as previously described (**Fig. 3**).^28^ The cameras were rigidly mounted above the Flexcell baseplate, to enable imaging of the top surface of the growth channel assembly, including the entire length of the channel. Each growth channel was imaged at 50 frames per second during cyclic mechanical strain, and non-contacting strain analysis was completed using VIC-3D 2009 digital image correlation software (Correlated Solutions, Inc.) to create an optical map of strains within the growth channel assembly. A virtual extensometer was then created along the length of the growth channel to calculate the tensile strain experienced within the growth channel. These measured tensile strains (output strains), averaged over the six independent growth channel assemblies, were plotted against strains programmed in Flexcell (input strains), and fit with a line of best fit to establish calibration curves.

**Figure 3.**
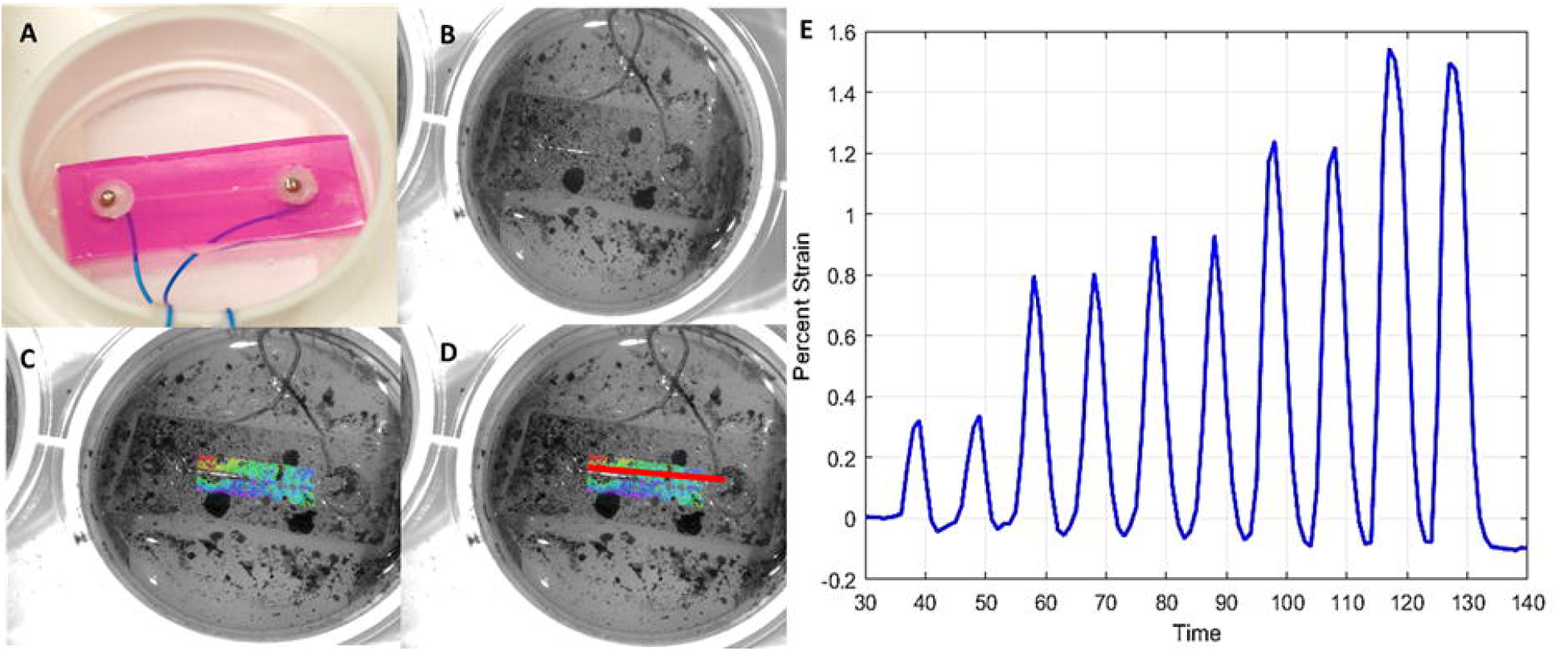
To characterize mechanical strains experienced by developing scaffold-free tissue constructs, (A) growth channel assemblies coupled to modified Dual Stimulation plates were (B) anisotropically speckled with a high-contrast pattern to (C) allow an optical strain map to be generated within the growth channel assembly using digital correlation (DIC). Creating (D) a virtual extensometer along the length of the growth channel allowed for measurement of applied sinusoidal cyclic strain within fiber growth channels. (E) Representative strains measured using DIC for growth channel assembly loaded for two cycles each at 0.50%, 0.70%, 0.75%, 1.0%, and 1.5% at 0.1Hz.

To test our ability to deliver precise tensile strains, we used the aforementioned calibration curves to determine the input magnitudes that should produce strains at each of the five desired strain magnitudes. We applied two cycles of each input strain magnitude (calculated from calibration curves) to six independent growth channel assemblies, at the three frequencies of interest. The resulting strains in the growth channel were measured dynamically using a DIC-based virtual extensometer, as described above.

#### 3.1.2. Electrical Characterization

Fabricated growth channel assemblies were coupled to Dual Stimulation Plates, as done for mechanical characterization, and individual plate wells were filled with DMEM (high conductivity, ∼1.5 S/m) to cover the growth channel. Wire leads from the Dual Stimulation plates were connected to a 16-Bit, 250kS/s M Series Multifunction Bus-powered Data Acquisition (DAQ) module (NI USB-6211, National Instruments, Austin, TX), and a LabVIEW Virtual Instrument was used to deliver a biphasic square waveform of desired magnitude, frequency, and duty cycle. For electrical stimulation characterization, biphasic pulses with amplitudes of 1, 5, or 8 V, designed to mimic neural impulses in muscle contraction, were applied to six independent growth channel assemblies at four different stimulation frequencies (0.2, 1.0, 2.0, and 10.0 Hz). The resulting voltages were measured via recording electrodes held at three distinct locations along the length of the growth channel - 0, 5, and 13 mm from the stimulating electrode - to assess the delivered electrical stimulation throughout the channel (**Fig. 4**). Average measured voltages at each frequency were plotted against voltages input in LabVIEW, and lines of best fit were applied to generate calibration curves.

**Figure 4.**
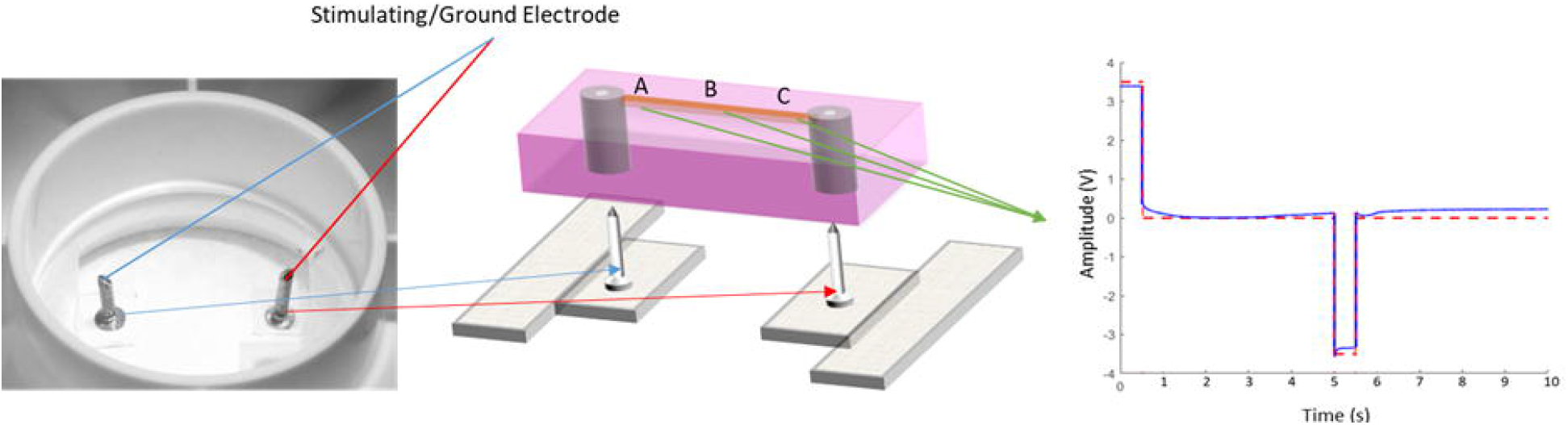
(Left) To incorporate electrical stimulation capabilities, Flexcell Tissue Train plates were further modified with soldered wires, such that the vertical loading pins doubled as stimulating electrodes. (Center) Electrical impulses applied via LabVIEW were measured using recording electrodes at three locations (positions A, B, and C) along the length of the growth channel. (Right) Representative time-voltage trace for a biphasic 3.5V square wave impulse (10% duty cycle) applied at 0.2Hz, showing controlled reproducible electrical stimulation of cells/cell-based constructs is possible using Dual Stimulation plates. (modified from [26]).

To test our ability to apply desired electrical stimulation, we applied voltages calculated using calibration curves to six independent growth channel assemblies at the three frequencies of interest, and recording electrodes placed along the length of the growth channel were used to record resulting electrical stimulation voltages.

#### 3.1.3. Combined Stimulation

Having calibrated mechanical and electrical stimulations independently, we assessed the potential effects of combined mechanical and electrical stimulation on independent calibrations. Six separate growth channels were sequentially subjected to: (i) independent mechanical stimulation, (ii) independent electrical stimulation, and (iii) concurrent electrical and mechanical stimulation using specific waveforms of interest for skeletal muscle tissue engineering.^27^ Growth channel assemblies were prepared for DIC strain assessment (*i*.*e*., blotted dry, speckle-coated), coupled to individual wells in Dual Stimulation Plates, then 2 mL of DMEM were added to the bottom of each well, with an additional 1 mL of DMEM pipetted over the growth channel.

First, we characterized independent mechanical stimulation, applying two cycles each of calibrated strain values for desired 0.50%, 0.70%, 0.75%, 1.00%, and 1.50% strains at 0.5 Hz to six independent growth channel assemblies, and measured the resulting uniaxial strain within the growth channels using DIC, as previously detailed. Next, we characterized independent electrical stimulation in the same six growth channel assemblies, using the established calibration curves to deliver a biphasic square pulse (1.5V, 10% duty cycle) at 1.0 Hz, and measured the resulting voltages. We measured the electrical measurements using a single recording electrode positioned at the growth channel’s center, as our calibration experiments showed voltage to be consistent along the growth channel length. Finally, we applied concurrent mechanical and electrical stimuli, identical to those applied independently, to the same growth channels used for independent characterization. We compared the resulting mechanical strains and stimulating voltages measured within the growth channel during concurrent stimulation to their respective independently-measured values to determine how combined stimulation affected independent calibrations.

#### 3.1.4. Statistical Analysis

A Single-Factor ANOVA was used to assess the differences between individual growth channel assemblies used for mechanical and electrical characterization. As no significant differences were observed across individual growth channels, in either mechanical or electrical tests (see results), two-tailed Student’s t-tests were used to examine differences between averaged output strains or voltages and the desired target values. A Two-Factor ANOVA was used to examine differences in mechanical or electrical stimulation when delivered independently or in combination. All data are reported as mean ± SD, with a significance level of α = 0.05 in all analyses.

### 3.2. EXPERIMENTAL RESULTS

#### 3.2.1. Mechanical Strain Characterization

Mechanical strains measured within the growth channels were smaller than their programmed input strains (0.50%, 0.70%, 0.75%, 1.0%, or 1.5%), indicating that the Flexcell plates’ modifications slightly attenuate the input strain. Additionally, plotting measured output strains against their corresponding input strains (**Fig. 5A**) revealed that mechanical calibration was frequency-specific, with slightly different calibration curves for each cyclic strain frequency (0.1, 0.5, 1 Hz). Data for the five strain amplitudes was well described by a linear calibration curve at each frequency - R^2^ = 0.845, 0.945, 0.941 - for oscillation frequencies of 0.1, 0.5, and 1 Hz, respectively (**Fig. 5A**).

**Figure 5.**
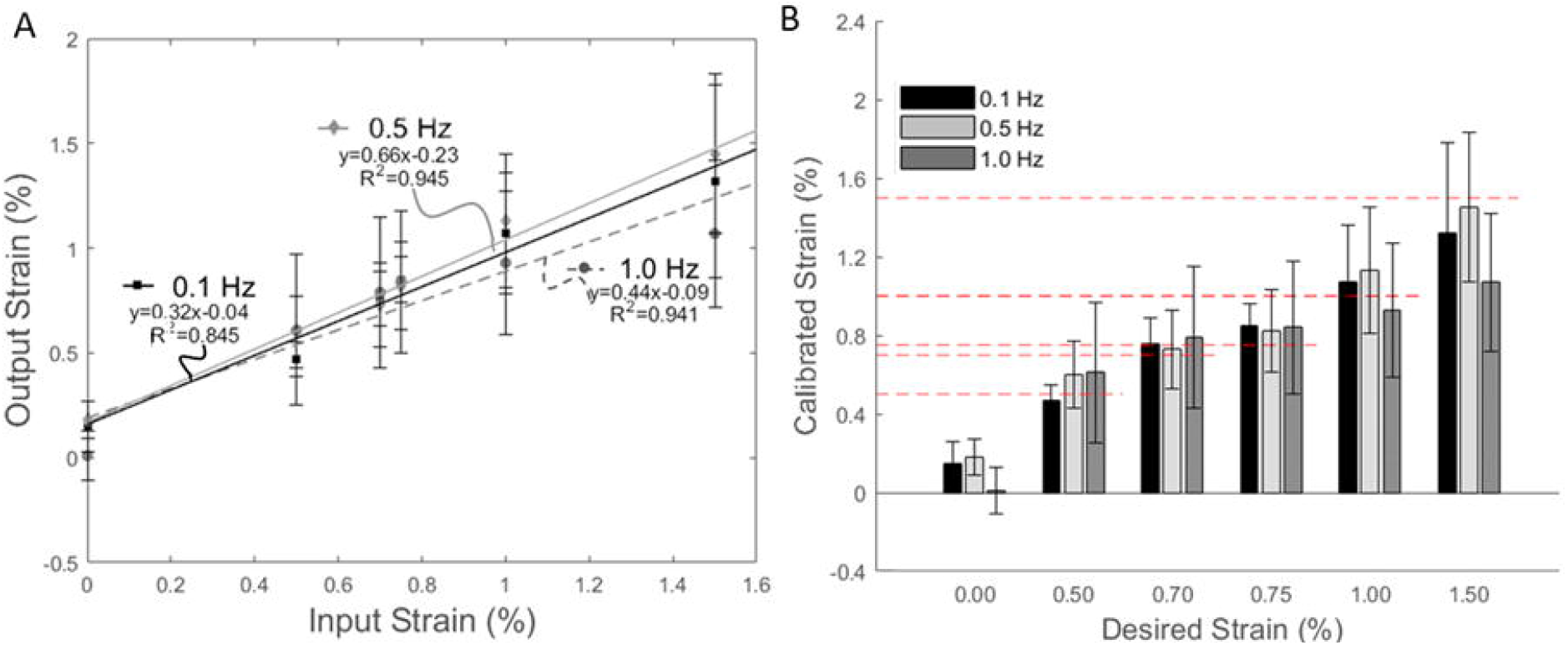
(A) Calibration curves at each frequency of mechanical loading were established by fitting DIC-measured output strain to input strain with a line of best fit. (B) By utilizing these established calibration curves to inform input strain, calculated input strain values produced calibrated strains within the growth channel that were not statistically different from their desired target strain values. Mean ± Standard deviation, n=6.

We used the calibration curves to determine the input strain values needed to produce desired output strains (*e*.*g*., to mechanically strain a fiber 0.5% at 0.5 Hz, 1.11% strain is programmed in the Flexcell computer). Calibration-determined input strains produced average output strains that were not statistically different from the intended target strains of 0.50%, 0.70%, 0.75%, 1.0%, or 1.5% (*p* > 0.05, **Fig. 5B**).

#### 3.2.2. Electrical Characterization

Programmed input voltages (1, 3, 8 V) were greater than their corresponding output values measured within the growth channel, at each location tested along the channel length. Voltages recorded at the three different locations were not statistically different within each channel; therefore, their average was used as the channel’s representative voltage measure. Measured output voltages showed a strong linear correlation (R^2^ = 0.99) with input voltages (**Fig. 6A**), and their values revealed that electrical calibration was not stimulation frequency-dependent.

**Figure 6.**
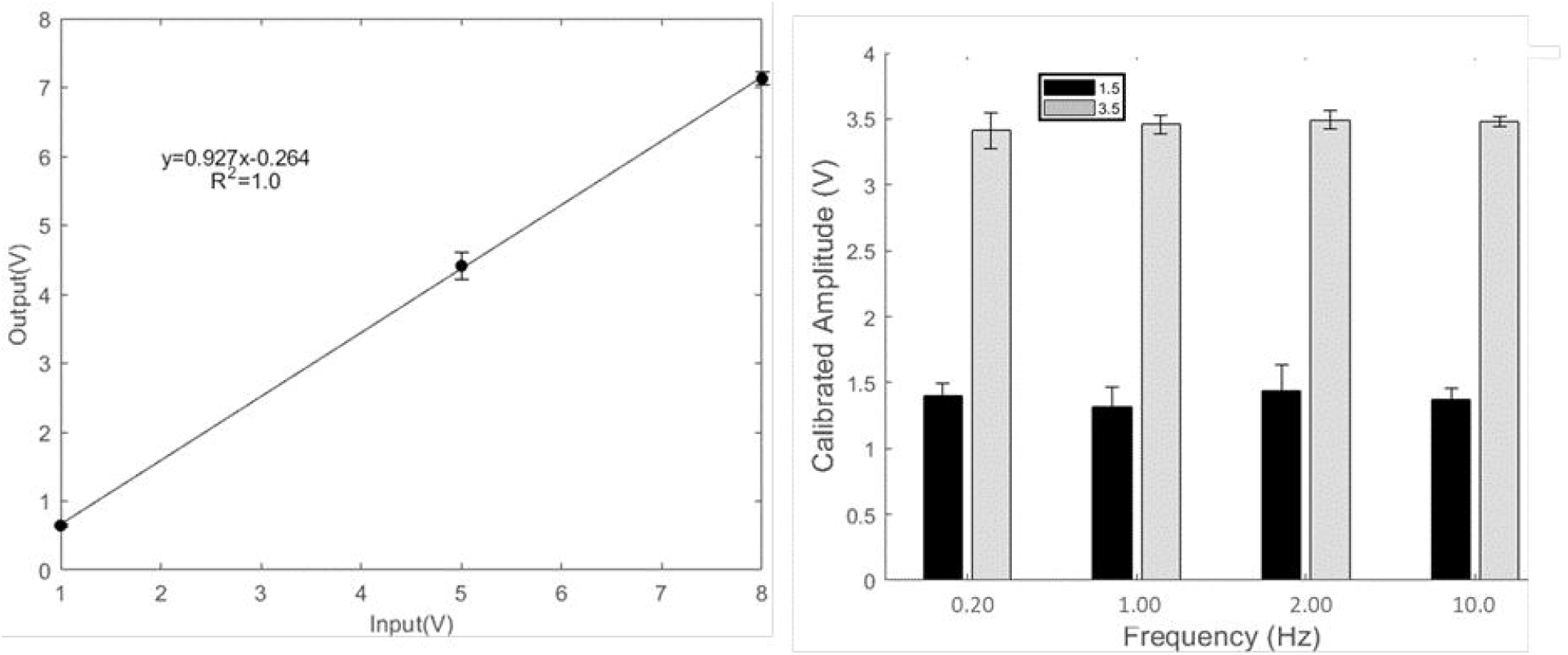
A) The electrical calibration curve, established by fitting recorded voltages across the different frequencies (0.2, 1, 2, and 10 Hz) and voltages (1, 5, and 8 V), was used to calculate input voltages required for desired output. B) Values achieved using the calibration curve for 1.5 Vs and 3.5 Vs were statistically similar to their desired target voltages, across all frequencies.

Similar to our approach for mechanical calibration, we used the calibration curve to calculate input voltages needed to produce desired output voltages (*e*.*g*., to electrically stimulate a fiber 1.5V at any frequency, 1.9V is programmed into LabVIEW). For the two voltages of interest (1.5V and 3.5V), the calibration-determined input voltages produced output voltages statistically similar (*p* = 0.032 for 1.5V and *p* = 0.022 for 3.5V) to the desired voltages at each frequency (**Fig. 6B**).

#### 3.2.3. Combined Stimulation

With the desire to apply concurrent mechanical and electrical stimulation, it was important to determine to what extent each stimulation affects the other’s individual calibration. The addition of electrical stimulation during concurrent stimulation had no significant effect on the independent mechanical strain stimulation calibration (< 2% difference and a −0.04 unloaded offset; **Fig. 7A**). Likewise, the electrical stimulation calibration was unaffected by the addition of cyclic mechanical strain (< 1% difference; **Fig. 7B**). Additionally, strains and voltages measured for concurrent mechanical and electrical stimulation were not statistically different from their desired target values. Taken together, these results demonstrate our bioreactor’s ability to deliver precise mechanical and electrical stimulation, alone or in combination, to developing scaffold-free tissue engineered fibers.

**Figure 7.**
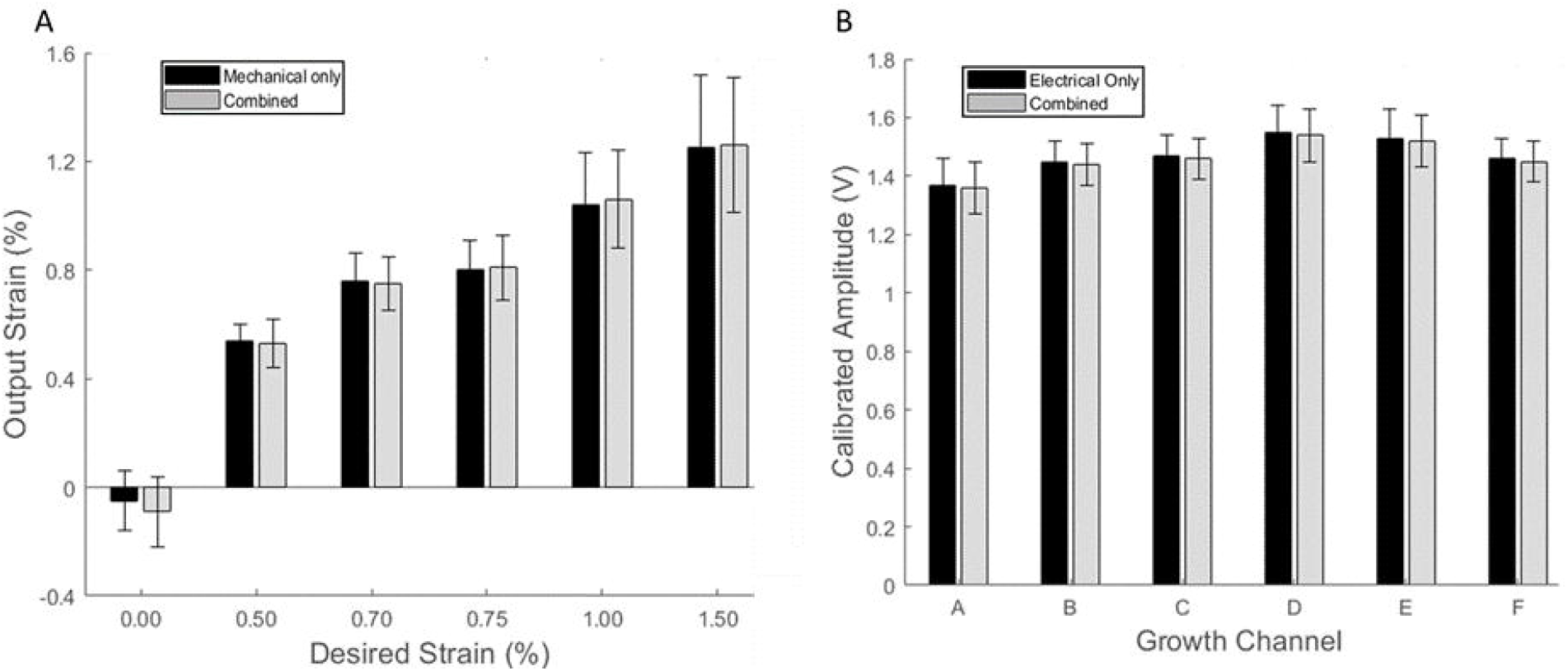
Concurrent delivery of mechanical and electrical stimulation had negligible impact on independent mechanical and electrical calibrations. A) There was no significant difference between mechanical strains measured during independent or concurrent stimulation. B) There was no difference in voltages measured during independent or concurrent stimulation for 6 independent growth channels. Thus, it appears that the mechanical stimulation is unaffected by concurrent electrical stimulation, and vice versa, allowing precise stimuli to be delivered either independently or in combination. Mean ± standard deviation; n=6.

## 5. DISCUSSION

It was critical to calibrate mechanical and electrical input signals to their output delivered within growth channels, to know the exact stimuli that the cells would receive. Our modifications to Flexcell plates altered their compliance and added mass, thereby affecting their factory calibrations. Prior to our calibration, the input mechanical strain magnitudes were consistently higher than the strains delivered to the growth channels. Additionally, these Dual Stimulation Plates (**Fig. 5**) exhibited different calibration curves than Flexcell plates used in our prior tendon engineering work, which were only modified with loading pins (*i*.*e*., no soldered wires; **Supplemental Fig. 1**). This difference is likely due to additional inertia and stiffness introduced by the wire leads. Therefore, our modifications for both mechanical and electrical stimulation affect how vacuum pressure applied by the Flexcell system translates to the tensile strains that cells experience within the growth channel. Because the mechanical strain is delivered by coupling growth channel assemblies to modified Dual Stimulation plates, any slight fabrication differences, in either the growth channels or the modified plates, would affect how the input strain is transferred into the growth channels, and their variances would be additive. Despite these variations, applied input strain magnitudes, determined using calibration curves, were not statistically different from their target desired strain values (0.50%, 0.70%, 0.75%, 1.0%, 1.5%), illustrating the ability to precisely deliver prescribed cyclic mechanical strain. We also observed that the mechanical calibration was frequency-dependent, with increased resistance to mechanical stimulation seen at higher frequencies. Yet, despite differing absolute values at the various frequencies (0.1, 0.5, 1.0 Hz), all calibrations were linear, reflecting the linear relationship between vacuum pressure and strain magnitude.

Electrical stimulation appeared to be frequency-independent, with a single line of best fit accurately representing all calibration frequencies (R^2^ = 0.99). Additionally, electrical measurements were very consistent, with no observed differences along the channel length, and very small channel-to-channel differences. Variance in electrical stimulation was far less than that of mechanical stimulation (*i*.*e*., smaller standard deviations); suggesting that measured voltages were much less affected by variations in plate or growth channel fabrication. Therefore, we believe that many of the variations that can affect mechanical strain (*e*.*g*., fabrication of plate, growth channel, plate/channel interface) have negligible influence on the electrical input signal delivery within the growth channel. As with mechanical stimulation, the input voltages applied using the calibration curve were statistically similar to the desired target values (1, 5, or 8V), highlighting our bioreactor’s capacity to provide controlled electrical stimulation.

Concurrent stimulation had a negligible influence on independent calibrations; causing a < 2% difference in the independent mechanical calibration, and < 1% difference in the independent electrical calibration. Thus, the mechanical and electrical calibrations remain valid, whether applied alone or in combination, thereby allowing concurrent stimulation and other more complex electrical/mechanical loading patterns to be explored. As a result, this bioreactor enables us to deliver precise, controlled, reproducible mechanical and electrical stimulations, alone or in combination, for tissue engineering applications.

An important novelty of this bioreactor is its capacity to apply prescribed stimuli to constructs *as they develop*. There is growing interest in the mechanobiology of tissue genesis and maturation, and scaffold-free tissue engineering approaches have emerged as attractive models to study tissue development. However, scaffold-free techniques have faced technical challenges in this area because the developing constructs need sufficient structural integrity to withstand applied mechanical stimulation. To overcome this, scaffold-free techniques typically allow the construct to develop significant structure before introducing mechanical loading. Applying stimulation after the construct has significantly formed promotes matrix remodeling, rather than tissue formation, and precludes investigation of how these stimuli influence tissue construct development.^7,21,31^ Our bioreactor can apply mechanical stimulation to scaffold-free tissue constructs as they develop, promoting matrix (re)generation.^26,27,31^

There is also great interest in the impact of electrical stimulation during engineered tissue development, particularly in electrically-active tissues, such as muscles and nerves. Electrical stimulation is suggested to occur throughout skeletal muscle development,^32,33^ and it has been exogenously applied to upregulate muscle specific genes, like pax-7,^32^ and increase muscle volume in aneurogenic muscles^33^ of chick embryos. Since electrical stimulation contributes to muscle development and biomechanical function, its use in *in vitro* tissue engineering could be beneficial, and ultimately translate to more functionally mimetic skeletal muscle constructs.^13^ Indeed, some studies have utilized electrical stimulation in skeletal muscle tissue engineering, and shown synergistic effects when delivered in combination with mechanical stimulation, as evidenced by increased matrix alignment, myotube formation, and force production.^6,20,34^ However, because those studies were conducted using scaffold-based methods, or by applying the stimuli to scaffold-free constructs after they had fully formed, they could not explore the effect of concurrent stimulation on construct development. Thus, this bioreactor’s ability to apply stimulation to scaffold-free tissue constructs as they develop can provide additional insight on the developmental and reparative roles of these stimuli, and their possible synergies, and may further enhance tissue engineering approaches.^7,13,21,31^

Our ongoing work in muscle tissue engineering aims to use this bioreactor to mimic key electrical and mechanical stimuli that promote myogenesis and engineer functional skeletal muscle fibers that possess similar passive properties to skeletal muscle and generate muscle-like specific force. We also seek to explore whether precise electrical and mechanical stimuli can promote fiber-type specification, *e*.*g*., applying higher frequencies of stimulation to promote faster fiber types, during development. Some of our preliminary findings, in engineered skeletal muscle fibers derived from C2C12 myoblasts, showed improved fiber longevity when subjected to combined mechanical and electrical stimulation, as compared to independent electrical stimulation or no stimulation.^30^ We will explore these potentially synergistic effects in future studies.

Although the calibrations explored in this study were for skeletal muscle tissue engineering, the bioreactor’s mechanical and electrical stimuli can be easily adjusted to accommodate a wide range of applications. In addition to the sinusoidal waveform assessed here, the modified Flexcell system allows square and sawtooth waveforms, at frequencies of 0.1 - 3 Hz, and strain magnitudes up to 20%, as well as simultaneous mechanical stimulation of up to 24 developing constructs. Similarly, the LabVIEW-DAQ module interface program is able to generate various waveforms, at different frequencies and voltages (± 10 V), and can deliver electrical stimulation to four developing constructs in its current configuration, with the capacity to be expanded for more. Thus, this inexpensive and high-throughput bioreactor allows for great flexibility in applied stimulation which may allow for studies of development of other tissues. However, stimulation at different magnitudes and frequencies than those explored herein may require further calibration to ensure that the programmed stimuli accurately translate to the cells.

In addition, the framework of this bioreactor allows it to be adapted to other platforms, including larger structural scales (*e*.*g*., fascicles, whole tissue), and scaffold-based constructs. While our immediate interest is in its utility for tendon and skeletal muscle fiber engineering, the bioreactor may be used to study the roles of mechanical and electrical stimulations in the development and maturation of other tissues, such as nerve, skin, ligament,^8^ smooth muscle,^35^ and cardiac muscle.^14^ This can be achieved by changing the cell source and adjusting the stimulation patterns to better mimic those relevant to each tissue’s development. Additionally, this bioreactor has potential for straightforward incorporation of chemical and environmental stimuli (*e*.*g*., growth factor addition, macromolecular crowding, hypoxia) to allow for controlled multimodal stimulation, alone or in combination. As the importance of biophysical stimulation in tissue engineering is recognized and incorporated, this bioreactor provides a platform to further our understanding of the distinct, and potentially synergistic, roles these stimuli play in tissue development and functional maturation, which may advance future tissue engineering approaches.

## 6. CONCLUSION

In this work, we created and robustly calibrated a bioreactor to deliver controlled, reproducible mechanical and electrical stimulation, independently or combined, to scaffold-free tissue-engineered constructs. We showed that once independent calibration curves were established, input parameters could be prescribed to apply specific cyclic mechanical strains (amplitude and frequency) and electrical stimulations (voltage, frequency, pulse width, waveform) to cells within fiber growth channels. Additionally, our independent calibrations were not affected by concurrent stimulation. Thus, this relatively simple bioreactor enables precise, dynamic mechanical and electrical stimuli to be delivered, either alone or in combination, to engineered scaffold-free fibers as they develop, to provide unique insight into the influence of biophysical stimuli on the development and functional maturation of engineered musculoskeletal tissues.

## ACKNOWLEDGMENTS

This work was supported, in part, by National Science Foundation (NSF) Career Award CBET-0954990 (DTC). Special thanks to Dr. Monica Agarwal for input on experimental design.

## AUTHOR DISCLOSURE STATEMENT

No competing financial interests exist.

---

**Supplemental Figure 1.** Calibration of plates for mechanical stimulation only. A) Calibration curves at each frequency of mechanical loading were established by fitting output strain measured using DIC for input strains programmed in Flexcell (0.50, 0.70, 0.75, 1.0, and 1.5%). (B) Calibrated strain values measured using DIC were not statistically different from Desired Strain values programmed using established calibration curves. Mean ± Standard deviation, n=6.

## Notes

### Competing Interest Statement

The authors have declared no competing interest.

