## Supplementary figures and images for "A bioreactor for controlled electromechanical stimulation of developing scaffold-free constructs"

### Supplemental Figure 1

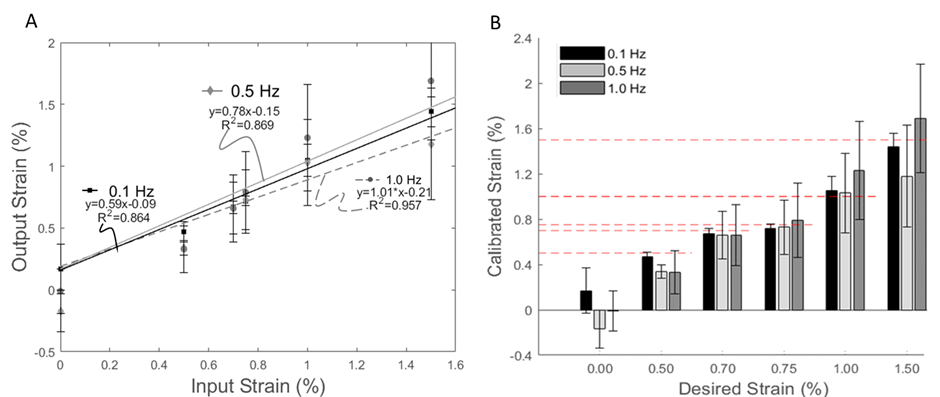
